# Hydrophobicity drives the systemic distribution of lipid-conjugated siRNAs via lipid transport pathways

**DOI:** 10.1101/288092

**Authors:** Maire F. Osborn, Andrew H. Coles, Annabelle Biscans, Reka A. Haraszti, Loic Roux, Sarah Davis, Socheata Ly, Dimas Echeverria, Matthew R. Hassler, Bruno M.D.C. Godinho, Mehran Nikan, Anastasia Khvorova

**Author notes:** These authors contributed equally to this work (Maire F. Osborn, Andrew H. Coles). Co-corresponding authors: Maire F. Osborn. Anastasia Khvorova.

## Abstract

Efficient delivery of therapeutic RNA is the fundamental obstacle preventing its clinical utility. Lipid conjugation improves plasma half-life, tissue accumulation, and cellular uptake of small interfering RNAs (siRNAs). However, the impact of conjugate structure and hydrophobicity on siRNA pharmacokinetics is unclear, impeding the design of clinically relevant lipid-siRNAs. Using a panel of biologically-occurring lipids, we show that lipid conjugation modulates siRNA hydrophobicity and governs spontaneous partitioning into distinct plasma lipoprotein classes *in vivo*. Lipoprotein binding influences siRNA distribution by delaying renal excretion and promoting uptake into lipoprotein receptor-enriched tissues. Lipid-siRNAs elicit mRNA silencing without causing toxicity in a tissue-specific manner. Lipid-siRNA internalization occurs independently of lipoprotein endocytosis, and is mediated by siRNA phosphorothioate modifications. Although biomimetic lipoprotein nanoparticles have been considered for the enhancement of siRNA delivery, our findings suggest that hydrophobic modifications can be leveraged to incorporate therapeutic siRNA into endogenous lipid transport pathways without the requirement for synthetic formulation.

## Introduction

For over a decade, the underlying obstacle preventing the widespread clinical use of small interfering RNA (siRNA)-based therapies has been efficient and safe *in vivo* delivery. siRNAs are large (~14 kDa), polyanionic macromolecules that require extensive modifications to improve their inherently-poor pharmacological properties (e.g. plasma half-life of <5 min before renal excretion)^1,2^. Lipid- and polymer-based nanoparticles, which are effective transfection agents *in vitro*, can prolong circulation time and improve stability and bioavailability *in vivo*. However, nanoparticle delivery is typically limited to clearance organs with fenestrated and/or discontinuous endothelium (e.g. liver, spleen, and certain tumors). Moreover, nanoparticles can interact with opsonin proteins that enhance clearance by macrophage phagocytosis.

Molecular-scale delivery of siRNAs with small targeting ligands, cell-penetrating peptides, or lipid conjugates may be a simple and effective alternative to nanocarrier-based methods. The most clinically-advanced siRNA conjugate, trivalent N-acetylgalactosamine (GalNAc)-siRNA, binds to the asialoglycoprotein receptor on hepatocytes with high selectivity, and triggers potent and durable (~6 months) gene silencing in patients^3,4^. The second major class of molecular siRNA conjugates are lipids, which have been shown to enhance circulation time and promote local and systemic delivery and efficacy. Cholesterol-modified siRNA, one of the first reported lipid conjugates, silences liver apolipoprotein B (ApoB) expression at high doses (50 mg kg^−1^), while α-tocopherol and fatty acid conjugates have also shown potential for liver delivery and gene silencing^5–7^. Productive internalization of cholesterol-conjugated siRNAs hinges on its interactions with circulating lipoproteins and tissue receptors^6^. However, the relationship between the chemical composition of lipid conjugates and siRNA pharmacology has not been characterized.

In order to define design parameters for optimizing therapeutic oligonucleotide delivery, we investigated the influence of structurally diverse lipids on conjugate-mediated siRNA biodistribution, efficacy, and safety *in vivo*. These studies were enabled by the use of a clinically-validated, chemically-modified siRNA scaffold that is nuclease resistant and metabolically stable^8^. Here, we demonstrate that lipid conjugation modulates the hydrophobicity of hsiRNA. We present evidence suggesting that oligonucleotide hydrophobicity governs pharmacokinetic behavior by driving selective, *in situ* incorporation into endogenous lipoprotein pathways. Lipid modification also enables potent siRNA-mediated mRNA silencing in lipoprotein receptor-enriched tissues, including liver, adrenal gland, ovary, and kidney. These data suggest that hydrophobicity likely directs the rank-order tissue distribution of lipidic oligonucleotide conjugates that may otherwise be intended to be specifically receptor- or cell-targeting.

## Results

### Design and synthesis of structurally-diverse lipid-hsiRNA conjugates

To evaluate the impact of lipid conjugation on the pharmacological properties of oligonucleotides, we synthesized a panel of structurally diverse conjugates using a hydrophobically-modified siRNA scaffold, termed hsiRNA (Fig. 1a, b)^9–11^. This scaffold consists of a 20-nucleotide (nt) antisense (guide) strand and a 15-nt sense (passenger) strand, and contains alternating 2′-*O*-methyl and 2′-fluoro sugar modifications^12^. The 5′-end of the antisense strand bears a terminal (E)-vinylphosphonate modification that increases siRNA tissue accumulation, extends the duration of silencing activity, and shields oligonucleotides from 5′-to-3′ exonucleases^13^. The terminal nucleotide backbones are fully phosphorothioated to further inhibit exonuclease-mediated degradation and to promote cellular internalization^14^. These extensive modifications are essential for evaluating conjugate-mediated delivery *in vivo* because partially-modified or unmodified siRNAs are rapidly degraded and cleared^8^.

**Figure 1.**
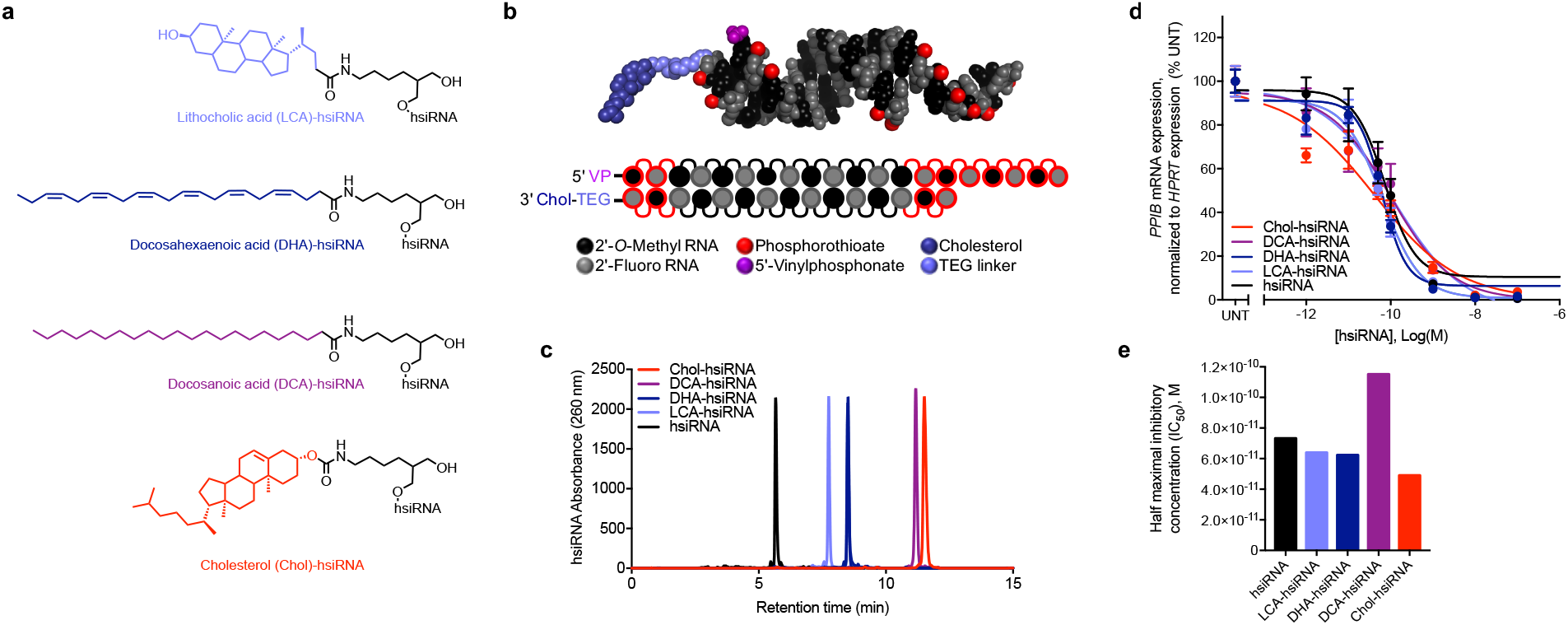
Synthesis and biophysical characterization of lipid-hsiRNA conjugates. (a) Chemical structures of lipid-hsiRNA conjugates. (b) Modification pattern and molecular model of lipid-hsiRNAs. (c) HPLC traces of lipid-hsiRNAs following reverse phase column chromatography. (d) HeLa cells were incubated with *PPIB*-targeting hsiRNAs at concentrations shown for 72 h. *PPIB* mRNA levels were measured using QuantiGene (Affymetrix), normalized to housekeeping *HPRT1* (hypoxanthine phosphoribosyltransferase 1) mRNA levels, and presented as percent of untreated control (n=3, mean +/− SD). UNT – untreated cells. (e) Plotted IC_50_ values determined from the best-fit curves in (d).

Previous studies have identified the 3′-end of the sense strand as an optimal position for conjugate attachment, with minimal effect on siRNA-RISC (intracellular RNA-induced silencing complex) loading^9,15^. Therefore, we designed cholesterol (Chol), lithocholic acid (LCA), docosahexaenoic acid (DHA), and docosanoic acid (DCA) conjugates to attach through a commercially available carbon-based linker to the 3′-end of the sense strand via an amide bond (Fig. 1a). Following standard solid-phase oligonucleotide synthesis and deprotection protocols (see **Methods**), hsiRNA conjugates were synthesized on a functionalized solid support bearing each individual lipid moiety. Synthesized oligonucleotides were then purified and characterized by high-performance liquid chromatography (HPLC) and liquid chromatography-mass spectrometry (LC-MS), respectively. All oligonucleotide sequences and chemical modification patterns used in this study are reported in **Supplementary Table S1**.

To evaluate the impact of lipid conjugation on hsiRNA hydrophobicity, we analyzed Chol-hsiRNA, DCA-hsiRNA, DHA-hsiRNA, LCA-hsiRNA, and unconjugated hsiRNA by reverse-phase HPLC and measured retention times (Fig 1c). Here, a longer retention time correlates with greater affinity for the hydrophobic (C8) stationary phase. Unconjugated hsiRNAs eluted relatively quickly (5.7 min), followed by LCA-hsiRNA (7.7 min), DHA-hsiRNA (8.5 min), DCA-hsiRNA (11.1 min), and Chol-hsiRNA (11.5 min). This result indicates that DCA- and Chol-hsiRNAs are more hydrophobic than DHA-, LCA-, or unconjugated hsiRNAs. Retention times did not directly correlate with the predicted partition coefficients of the free lipids (**Supplementary Table S2**), suggesting that lipid orientation and conjugation position influence overall siRNA hydrophobicity.

To assess the effect of lipid conjugation on RISC loading and siRNA silencing efficiency *in vitro*, we transfected each lipid-hsiRNA into HeLa cells and measured mRNA levels of a well-validated housekeeping gene, cyclophilin B (*PPIB*) (Fig. 1d). Calculated half-maximal inhibitory concentrations (IC_50_s) ranged from 4.9×10^−11^ to 1.2×10e^−10^ M, confirming that conjugation at the 3′-end of the sense strand does not substantially affect RNAi potency (Fig.1e). In summary, we successfully synthesized a panel of lipid-hsiRNA conjugates with varying degrees of hydrophobicity that retain gene silencing activity *in vitro*.

### Lipid conjugation reduces hsiRNA kidney exposure and promotes broad biodistribution

To evaluate the biodistribution of each lipid-hsiRNA conjugate *in vivo*, we administered Cy3-labeled lipid-hsiRNAs into mice by a single, subcutaneous injection (n = 2, 20 mg kg^−1^) and measured fluorescence distribution after 48 h. As lipid-hsiRNA hydrophobicity increased, we observed a clear, progressive reduction in kidney accumulation and an increase in liver retention (Fig. 2a). We confirmed this result using a peptide-nucleic acid (PNA) hybridization assay, which measures hsiRNA antisense strand concentration in tissue lysate by HPLC. Unconjugated hsiRNAs were subject to acute renal clearance after administration and reached peak tissue concentrations of ~1,300 ng hsiRNA/mg kidney (Fig. 2b). As lipid-hsiRNA hydrophobicity increased, kidney accumulation decreased over 18-fold (peak concentration of ~70 ng hsiRNA/mg kidney for Chol-hsiRNA). Our findings agree with previous pharmacokinetic analyses of lipid-conjugated hsiRNAs, which found that the initial distribution half-life of Chol-hsiRNA was 33-35 min, but was only 15-17 minutes for unconjugated hsiRNA^16^. Moreover, the total exposure over time (area under the curve; AUC) increased >10-fold for Chol-hsiRNA compared to unconjugated hsiRNA (unpublished data)^16^. Unlike the kidney, oligonucleotide concentrations in the liver increased with increasing hydrophobicity of the lipid conjugate (~75 ng mg^−1^ for unconjugated hsiRNA up to ~715 ng mg^−1^ for Chol-hsiRNA). Lipid conjugation also enabled delivery beyond the liver and kidney, supporting conjugate-mediated siRNA accumulation in lung, heart, adrenal gland, spleen, adipose tissue, and femoral muscle (Fig. 2b).

**Figure 2.**
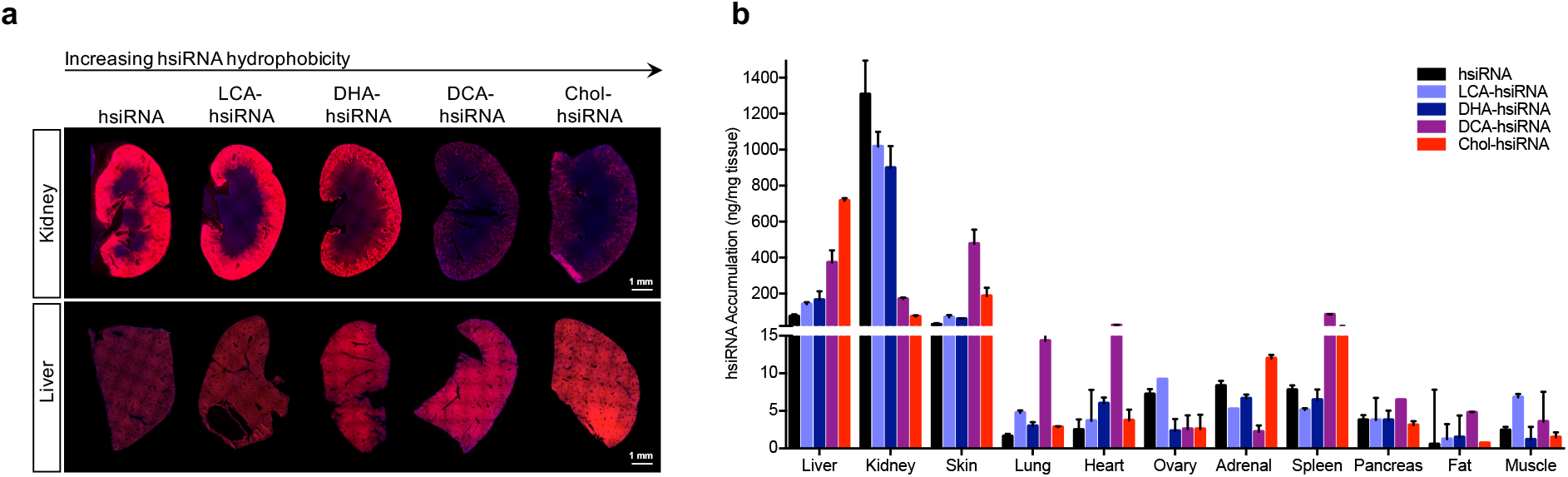
Systemic biodistribution and tissue accumulation of lipid-hsiRNA conjugates. Biodistribution of lipid-hsiRNAs 48 h after a single, subcutaneous injection (n = 3 mice, 20 mg kg^−1^) (a) Kidney and liver distribution of lipid-conjugated hsiRNAs. Cy3-labeled lipid-hsiRNAs (red), nuclei stained with DAPI (blue). (b) Guide strand quantification of Cy3-labeled lipid-hsiRNAs by a PNA hybridization-based assay. Data presented as mean ± SD.

### Hydrophobicity governs the interaction between lipid-hsiRNAs and serum proteins

To investigate potential mechanism(s) governing the association between lipid-siRNA hydrophobicity and biodistribution, we measured the affinity of lipid-siRNAs toward plasma constituents both *in vitro* and *in vivo*. Binding to albumin, a permissive host for both endogenous and exogenous ligands^17^, has been proposed to improve the pharmacokinetics of lipid-conjugated oligonucleotides ^7^. Thus, we measured the affinity of Cy3-labeled lipid-hsiRNAs and bovine serum albumin (BSA) *in vitro* using native gel electrophoresis (**Supplementary Fig. S1**). Unmodified hsiRNA did not associate with BSA to any measureable degree. However, all lipid-hsiRNA conjugates formed stable complexes with BSA, and exhibited dose-dependent migration shifts with increasing albumin concentration. We observed a direct correlation between hsiRNA hydrophobicity and BSA binding, with DCA-hsiRNA and Chol-hsiRNA exhibiting the highest affinities (K_d_ ~ 120 and 164 µM, respectively).

To more precisely characterize the behavior of lipid-conjugated siRNAs in the circulation, we used high-resolution size exclusion chromatography (SEC) to measure the interaction between oligonucleotides and circulating proteins in mouse serum. Previous reports have suggested that sterol-, bile salt-, and fatty acid-siRNAs are capable of associating with both albumin and lipoproteins in the bloodstream^6^. Thus, we used SEC to define relative size zones associated with very low-density lipoprotein (VLDL), low-density lipoprotein (LDL), and high-density lipoprotein (HDL) particles, in addition to small proteins (predominantly albumin) from mouse serum (Fig. 3a). Peak assignments were confirmed by measuring cholesterol content in isolated fractions (**Supplementary Fig. S2**). Assessment of the serum SEC profiles of wild-type (FVB/NJ) male and female animals (**Supplementary Fig. S2**) revealed no apparent sex-dependent differences in lipoprotein peak assignment. Therefore, we performed subsequent analyses in female animals only.

**Figure 3.**
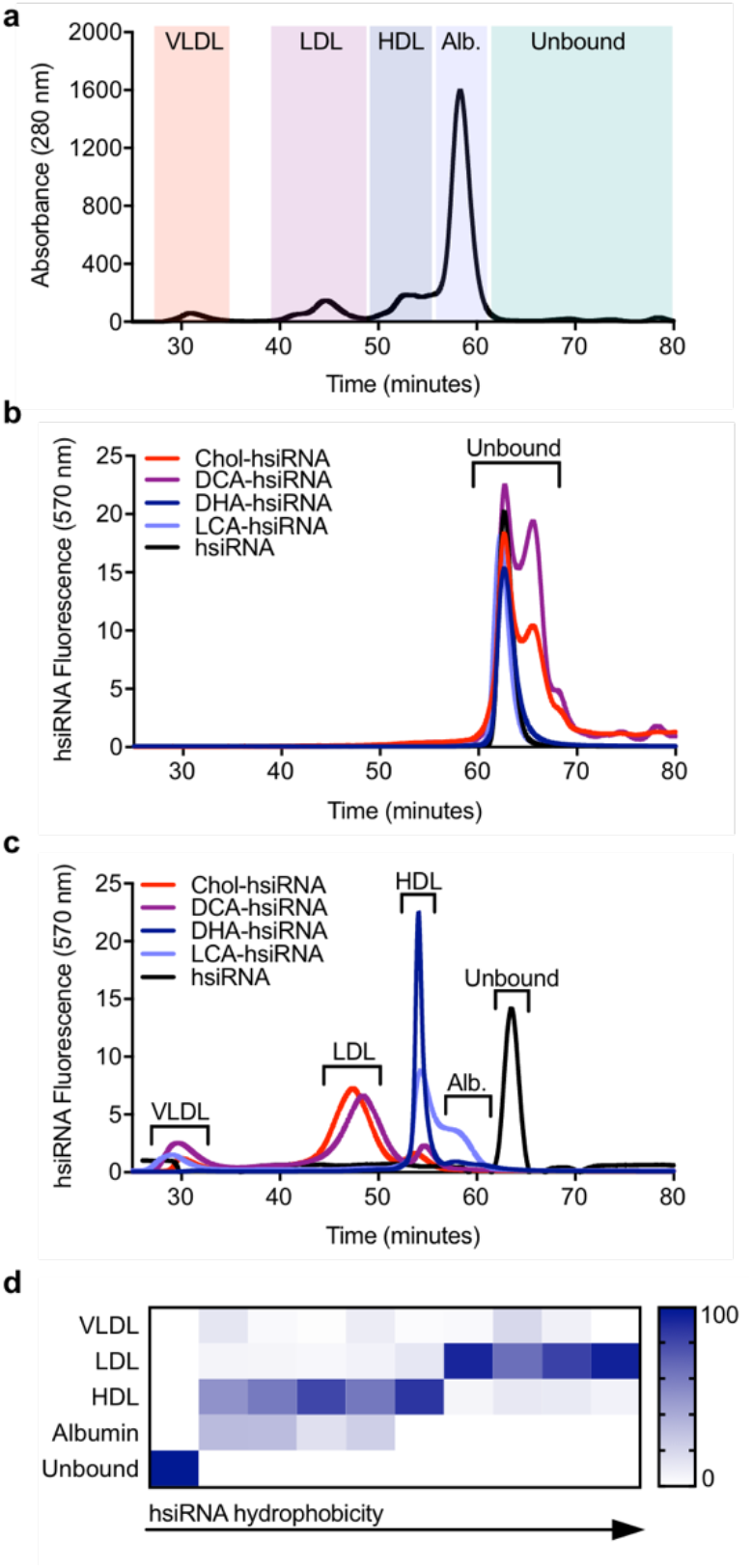
Lipoprotein binding profiles of lipid-conjugated hsiRNAs. (a) Mouse serum protein distribution following size exclusion chromatography (SEC). Red shading: VLDL; purple shading: LDL; dark-blue shading: HDL; light-blue shading: albumin; green shading: no protein. (b) Retention times of Cy3-labeled lipidhsiRNAs following SEC. (c) Average retention times of Cy3-labeled lipid-hsiRNAs in mouse serum, 15 minutes after IV injection (n = 2). Peak shifts indicate serum protein association. (d) Summary of peak integrations of lipoprotein binding profiles for a variety of lipid-hsiRNAs.

To determine if lipid-hsiRNAs form stable complexes with lipoprotein particles or albumin *in vivo*, we first measured baseline retention times of free Cy3-labeled unconjugated and lipid conjugated-hsiRNAs (Fig. 3b). The unbound forms of unconjugated hsiRNA, LCA-hsiRNA, and DHA-hsiRNA eluted at 62.9 min as a single peak. Unbound DCA-hsiRNA and Chol-hsiRNA eluted as a doublet, with peaks at both 62.9 min and 65.8 min. The presence of a doublet may reflect a propensity for these two hydrophobic species to form reversible macromolecular structures under non-denaturing conditions. Next, we analyzed serum samples from mice injected intravenously with Cy3-labeled Chol-siRNA, DCA-siRNA, DHA-siRNA, LCA-siRNA, or unconjugated siRNA (n=2, 20 mg kg^−1^). Given the average elimination half-lives of Chol-hsiRNA and unconjugated hsiRNA^16^, we collected serum 15 minutes after injection. By monitoring the baseline shift in hsiRNA fluorescence signal, we determined that each lipid conjugate has a unique lipoprotein binding signature *in vivo* (Fig. 3c). In direct contrast to our *in vitro* results, we discovered that lipid-hsiRNA conjugates do not significantly associate with albumin in serum, likely due to the presence of higher affinity chaperones, such as lipoproteins. Unconjugated hsiRNAs did not associate with any serum components and were rapidly excreted into the urine (Fig 2a, Fig. 3c). LCA-hsiRNA associated predominantly with HDL (50.4% of total binding) and albumin (32.2%), with residual binding towards VLDL (12.2%) and LDL (5.3%). DHA-hsiRNA exhibited a stronger affinity for HDL (80.5%), with lesser co-migration with albumin (14.9%), LDL (3.7%), and VLDL (1%) peaks. Conversely, DCA-hsiRNA and Chol-hsiRNA were predominantly bound to LDL (65% and 82%, respectively), with residual binding to VLDL and HDL. Peak integrations (Fig. 3c) are summarized in **Supplementary Table S3**.

To further investigate the similarity between the lipoprotein binding profiles and degree of hydrophobicity of siRNAs, we employed the same SEC methodology to analyze an expanded panel of lipid-hsiRNA conjugates. As described previously, we performed SEC-based profiling of lipoprotein-hsiRNA binding and plotted these data as a function of relative lipid-hsiRNA hydrophobicity (Fig. 3d; **Supplementary Table S3**; Biscans *et al.*, submitted). We observed a strong correlation between hsiRNA hydrophobicity and segregation into albumin, HDL, and LDL peaks. Our findings suggest that oligonucleotide hydrophobicity can be engineered to achieve predictable and stepwise partitioning into different lipid transport pathways *in vivo*.

### Lipid-hsiRNAs elicit sustained systemic *in vivo* gene silencing after a single injection

Given that lipid-hsiRNAs predominantly complexed with circulating lipoproteins after systemic administration, we assessed the cell-specific distribution and gene-silencing efficacy of lipid-hsiRNAs in tissues that are enriched in lipoprotein receptors. For all tissues, cell-specific distribution of hsiRNAs was measured 48 h after a single, subcutaneous injection into mice (n = 3, 20 mg kg^−1^). Gene-silencing efficacy was determined by quantifying *Ppib* mRNA levels one week after a single, subcutaneous injection of *Ppib*-targeting hsiRNAs (n = 6, 20 mg kg^−1^).

#### Liver

LDL is the principal intercellular transporter of cholesterol, and LDL receptors (LDLR) are abundantly expressed on liver hepatocytes, adrenal cortex, bronchial epithelial cells, and adipocytes. We first monitored cell tropism in the liver and found that LDL-associated oligonucleotides (Chol- and DCA-hsiRNA) were readily visualized within cytoplasmic foci in liver hepatocytes (Fig. 4a, open arrows), while HDL- and albumin-associated hsiRNAs (DHA-, LCA-, and unconjugated hsiRNA) were undetectable (Fig. 4a, open arrows). By contrast, we did observe oligonucleotide internalization in Kupffer cells (stellate macrophages) for all five hsiRNAs (Fig. 4a, closed arrows). This result is consistent with what is known regarding the mononuclear phagocyte (reticuloendothelial) system as a first response against exogenous substances, including siRNAs and antisense oligonucleotides. We confirmed our findings for Chol-hsiRNA, DCA-hsiRNA, and DHA-hsiRNA by FACS, noting hsiRNA accumulation in 28-59% percent of endothelial cells and 18-49% of Kupffer cells, respectively (n = 3, 10 mg kg^−1^, **Supplementary Fig. S3**). Next, we correlated hsiRNA accumulation with *Ppib* mRNA levels. As expected, Chol-hsiRNA and DCA-hsiRNA displayed the highest activity, reducing liver *Ppib* mRNA expression by 59% and 65%, respectively (Fig. 4b). Unconjugated hsiRNA reduced *Ppib* mRNA levels by 39% (Fig. 4b). We did not observe a reduction in *Ppib* mRNA after administration of non-targeting control hsiRNAs (n = 6, 20 mg kg^−1^, **Supplementary Fig. S4**). We also did not observe significant upregulation of liver toxicity biomarkers, alkaline phosphatase (ALP) and alanine aminotransferase (ALT), in the blood (**Supplementary Fig. S5**).

**Figure 4.**
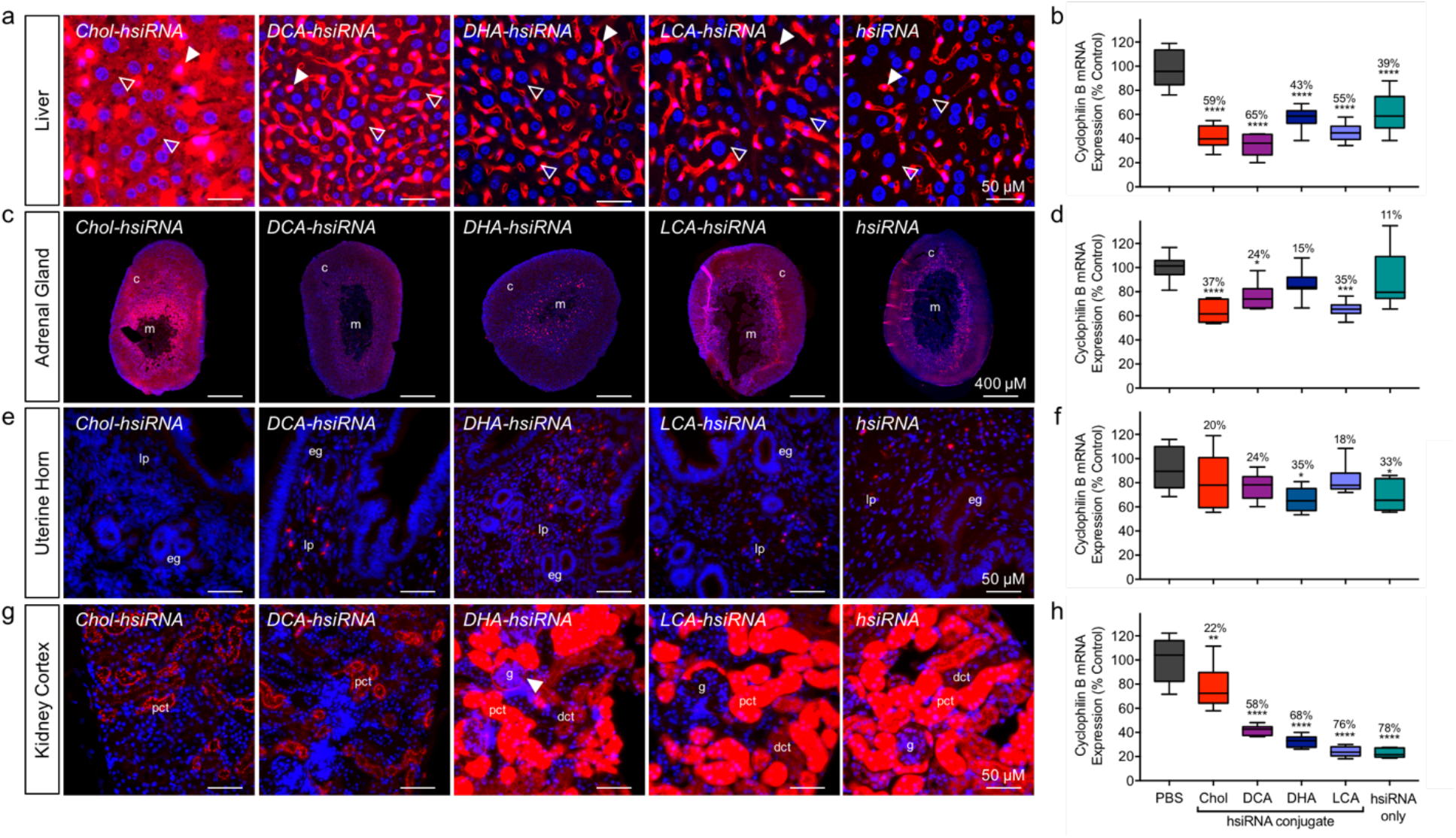
Distinct cellular uptake and efficacy patterns of lipid-conjugated hsiRNAs. Tissue-dependent internalization of lipid-hsiRNAs 48 h after a single, subcutaneous injection (n = 3 mice, 20 mg kg^−1^) in (a) liver, (c) adrenal gland, (e) uterine horn, and (g) kidney cortex. Cy3-labeled lipid-hsiRNAs (red), nuclei stained with DAPI (blue). Arrowheads described in text. c: cortex; m: medulla; lp: lamina propria; eg: endometrial gland; pct: proximal convoluted tubule; dct: distal convoluted tubule; g: glomerulus. Quantification of *Ppib* silencing by non-labeled lipid-hsiRNAs in (b) liver, (d) adrenal gland, (f) uterine horn, and (h) kidney cortex. *Ppib* mRNA levels were measured with QuantiGene 2.0 (Affymetrix) assay and normalized to a housekeeping gene, *Hprt*. All data presented as percent of saline-treated control. All error bars represent mean ± SD. *P < 0.05; **P < 0.01; ***P < 0.001; ****P < 0.0001 as calculated by one-way ANOVA with Tukey’s test for multiple comparisons.

#### Adrenal Gland

Circulating lipoproteins contribute over half of the cholesterol required for steroidogenesis in adrenocortical cells^18^. Therefore, we wanted to assess the pattern of lipid-hsiRNA distribution within the adrenal gland. As a general trend, higher levels of lipid-hsiRNA fluorescence were observed in the adrenal cortex compared to the medulla (Fig. 4c). The sterol-conjugated hsiRNAs (Chol and LCA) presented the highest levels of internalization in the adrenal cortex (Fig. 4c), and the strongest activity. Chol-hsiRNA silenced *Ppib* mRNA up to 37%, while LCA-hsiRNA reduced *Ppib* mRNA levels by 35% (Fig. 4d). DCA-, DHA-, and unconjugated hsiRNA showed lower levels of internal fluorescence in the adrenal cortex, and induced 24%, 15%, and 11% *Ppib* silencing, respectively (Fig. 4d).

#### Uterus

HDL is an integral component of the reverse cholesterol transport pathway, which transports lipids from extrahepatic tissues to the liver for excretion or recycling. Lipid transfer is commonly mediated by scavenger receptor B1 (SR-BI), which is highly expressed in liver sinusoidal endothelial cells, adrenal gland, ovary, and testis. Therefore, we monitored lipid-hsiRNA accumulation and activity within the uterus of female mice. DCA-, DHA-, LCA-, and unconjugated hsiRNAs showed punctate staining throughout the lamina propria, with minimal staining of the endometrial glands (Fig. 4e). We measured significant levels of *Ppib* silencing for both DHA-hsiRNA (35%) and unconjugated hsiRNA (33%) in biopsies from the uterine horn (Fig. 4f).

#### Kidneys

LCA-, DHA-, and unconjugated hsiRNAs were distributed similarly within the kidney cortex, staining both proximal and distal convoluted tubules (Fig. 4g). Minor levels of fluorescence in the glomerulus were detected for all three conjugates, with DHA-hsiRNA showing the most profound staining within the Bowman’s capsule (Fig. 4g, closed arrow). Chol-hsiRNA and DCA-hsiRNA were detected primarily within proximal convoluted tubules (Fig. 4g). After measuring *Ppib* mRNA levels in biopsies from the kidney cortex, we noted parallels between accumulation and activity, with unconjugated hsiRNAs showing the highest level of silencing (78%) (Fig. 4h). We did not observe any variability in biomarkers of kidney toxicity, including blood urea nitrogen and creatinine levels (**Supplementary Fig. S5**). We also monitored other standard serum toxicological markers, including electrolytes (Ca^2+^, phosphate, Na^+^, K^+^), albumin, globulin, bilirubin, and glucose, and saw no significant changes in total concentration after hsiRNA, DHA-hsiRNA, or DCA-hsiRNA treatment (**Supplementary Fig. S5**).

Taken together, these data suggest that lipid-conjugated hsiRNAs engage distinct lipid transport pathways and elicit gene silencing in a variety of tissues with minimal systemic toxicity. Interestingly, in some tissues, we observed a non-linear relationship between hsiRNA tissue accumulation and silencing efficacy, suggesting the presence of both productive and non-productive internalization pathways. This has been described for phosphorothioate-modified antisense oligonucleotides, which are intrinsically more active in hepatocytes than in the nonparenchymal cells of the liver^19^. The observed discrepancy between levels of hepatocyte internalization after 48 h and mRNA silencing after 1 week may indicate translocation of hsiRNAs from sinusoidal depots into hepatocytes, or the presence of inherently more potent endocytic pathways in hepatocytes (Fig. 4a, b). Whether this extends to the kidney or other tissues is unknown.

### hsiRNAs are internalized independently of LDL recycling via the LDL receptor

To probe the dependency of hsiRNA internalization on lipoprotein receptor expression, we compared the systemic biodistribution of an LDL-associated hsiRNA (DCA-hsiRNA) and an HDL-associated hsiRNA (DHA-hsiRNA) in both wild-type and LDLR-deficient mice. We hypothesized that if LDL-associated hsiRNAs are dependent on LDL endocytosis for internalization, their biodistribution should be significantly perturbed by LDLR depletion. LDLR-dependent internalization into the liver has been described for both cholesterol- and α-tocopherol-conjugated duplex siRNAs^6,20^. We intravenously injected wild-type (C57BL/6J) and LDLR-deficient (LDLR^−/−^) mice with DHA- or DCA-hsiRNA and quantified tissue accumulation by PNA hybridization (n = 3, 10 mg kg^−1^). As expected, DHA-hsiRNA systemic distribution and tissue internalization remained unchanged between wild-type and LDLR^−/−^ animals (Fig. 5a, blue bars). Unexpectedly, DCA-hsiRNA liver accumulation increased two-fold in the absence of LDLR (Fig. 5a, purple bars). This finding suggests that LDL-associated oligonucleotides are internalized in the liver by a mechanism independent of lipoprotein endocytosis. To determine if the observed increase in liver accumulation was cell-type specific, we acquired high-resolution images of the liver following subcutaneous administration of Cy3-labeled DCA-hsiRNAs (n = 2, 10 mg kg^−1^). Hepatocyte fluorescence in LDLR^−/−^ mice was visibly increased compared to wild-type mice (Fig. 5b). Image quantification revealed a two-fold increase in both cytoplasmic and nuclear fluorescence in mutant animals (Fig. 5c). Taken together, these data suggest that hepatocyte internalization of LDL-associated hsiRNAs can occur independently of LDL endocytosis through LDLR.

**Figure 5.**
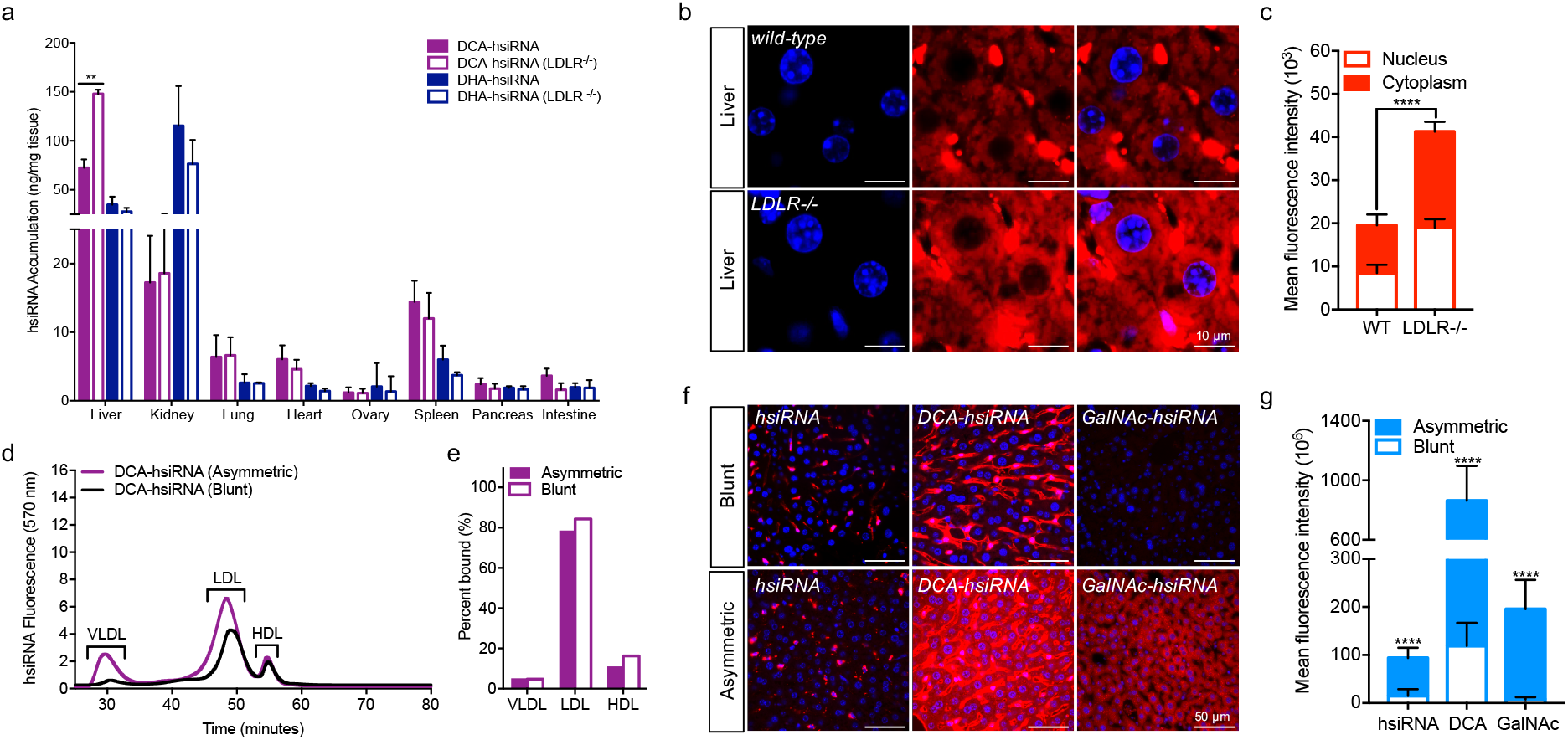
Mechanistic analysis of lipid-hsiRNA internalization in liver. (a) Guide strand quantification of Cy3-labeled DHAhsiRNAs and DCA-hsiRNAs in wild-type (C57BL/6J) and LDLR deficient animals after a single, intravenous injection (n = 3 mice, 10 mg kg^−1^) using a PNA hybridization-based assay. Data presented as mean ± SD. (b) Hepatocyte internalization of Cy3-labeled DCA-hsiRNA in wild-type and LDLR deficient animals after a single, intravenous injection (n = 3 mice, 10 mg kg^−1^). Image is representative. Cy3-labeled DCA-hsiRNAs (red), nuclei stained with DAPI (blue). (c) Quantification of fluorescent signal from images acquired in (b). (d) Average retention times of Cy3-labeled DCA-hsiRNAs in mouse serum, 15 minutes after IV injection (n = 2 mice, wild type or LDLR^−/−^). (e) Average peak integrations from lipoprotein profiles in (d). (f) Hepatocyte internalization of Cy3-labeled blunt and asymmetric siRNAs (unconjugated, DCA-conjugated, or GalNAc-conjugated) after a single, subcutaneous injection (n = 3 mice, 20 mg kg^−1^), staining as described in (b). (g) Quantification of fluorescent signal from images acquired in (f).

### hsiRNA hepatocyte internalization is facilitated by a phosphorothioate-modified, single-stranded overhang

A defining feature of the hsiRNA constructs used in this study is the presence of a 5-nt single-stranded, phosphorothioate-modified (PS) overhang, which resembles the fully PS, single-stranded backbone of conventional antisense oligonucleotides (ASOs). Recent studies investigating ASO internalization by hepatocytes have established that the asialoglycoprotein receptor (ASGPR) contributes to the uptake of unconjugated PS ASOs^21,22^. Therefore, one of the hepatocyte uptake pathways for DCA-hsiRNA may be a receptor-mediated process that requires a single-stranded, PS-modified overhang. To probe the structural requirements for hepatocyte uptake, we compared the asymmetric DCA-hsiRNA construct used throughout this study with a 20-nt blunt-ended DCA-conjugated hsiRNA. After analyzing the lipoprotein binding profiles of the asymmetric and blunt DCA-hsiRNAs, we observed no differences in lipoprotein binding character (Fig. 5d). Similar to DCA-hsiRNA, blunt DCA-hsiRNA primarily associated with LDL (84.3%) and exhibited less affinity for HDL (11.1%) and VLDL (4.8%) (Fig. 5e). This result provides further confirmation that lipoprotein association is mediated by the lipid conjugate and suggests that these compounds have comparable elimination half-lives *in vivo*. However, following a single, subcutaneous injection of Cy3-labeled blunt or asymmetric DCA-hsiRNA in wild-type mice (n = 2, 20 mg kg^−1^), we observed significantly less fluorescent signal in hepatocytes for blunt DCA-hsiRNA (Fig. 5f). This suggests that the single-stranded PS overhang contributes significantly to hepatocyte uptake. To determine whether the presence of a single-stranded PS region is sufficient for unconjugated hsiRNA internalization, we compared the liver distribution pattern of blunt and asymmetric unconjugated hsiRNAs following administration to mice (n = 2, 20 mg.kg^−1^). Although liver exposure of unconjugated hsiRNAs is limited, due to its rapid elimination half-life, we still observed a significant increase in hepatocyte fluorescence of asymmetric compared to blunt hsiRNA (Fig. 5f, g). To further validate our finding that single-stranded PS modifications enhance hepatocyte uptake, we synthesized triantennary GalNAc-conjugated blunt and asymmetric hsiRNAs. GalNAc-conjugated siRNAs are specifically and rapidly internalized in hepatocytes through the ASGPR receptor. Here, we detected a marked increase in GalNAc-mediated hepatocyte internalization in the presence of a single-stranded PS tail (Fig. 5f, g). It is unclear whether this is due to an increased affinity for ASGPR or mediated by a separate receptor.

## Discussion

We present the following model for the systemic distribution of lipid-conjugated, chemically-stabilized hsiRNA: After intravenous or subcutaneous administration, unconjugated hsiRNAs are rapidly filtered in the kidneys from the blood into the glomerular space, where they are reabsorbed (likely via scavenger receptors) by proximal tubule cells or excreted through the urine. This pharmacokinetic behavior has been observed for both blunt and asymmetric unconjugated siRNAs^23,24^. The use of an advanced chemical modification pattern affords enhanced metabolic stability^8^, and despite their short residence time, unconjugated hsiRNAs achieve ~80% target mRNA silencing in the kidney, providing a therapeutic strategy for kidney indications^25^. Conversely, lipid-conjugated hsiRNAs spontaneously associate with circulating lipoproteins, which increases plasma half-life and promotes exposure to extrarenal tissues. The lipid transport pathway engaged by lipid-conjugated hsiRNAs may be determined *a priori* following measurement of lipid-siRNA hydrophobicity. More hydrophobic hsiRNAs preferentially bind LDL, while relatively less hydrophobic hsiRNAs associate more with HDL. LDL-associated hsiRNAs enter into the endogenous lipid transport pathway and are preferentially internalized in LDL receptor-enriched tissues, such as liver, lung, and intestine. HDL-associated hsiRNAs are transported via reverse cholesterol transport and exhibit enhanced uptake and efficacy in SR-BI-enriched tissues, such as adrenal gland and ovary. hsiRNA internalization in the liver appears to occur independently of lipoprotein endocytosis, as LDL-associated hsiRNA are efficiently internalized into the hepatocytes of LDLR-deficient animals. This internalization is likely mediated by the single-stranded PS overhang, in a mechanism similar to that of ASOs^21,22^. Our theory is substantiated by the observation that hsiRNAs containing a PS tail are endocytosed by hepatocytes *in vivo* at significantly higher levels than blunt hsiRNAs, even when conjugated to a triantennary GalNAc ligand.

Extracellular lipoprotein trafficking of native small RNAs has been described by others ^26–28^. For example, microRNAs (miRNA) are known to be packaged into HDL and functionally transported through the circulation to recipient cells^27–29^, and biophysical studies suggest that nucleic acid-lipoprotein binding is mediated by divalent cation bridging to phosphocholine head groups^30–34^. However, we did not observe *in situ* assembly of unconjugated hsiRNAs with lipoproteins in mouse serum, suggesting that an entropic driving force, such as a lipid anchor, is required for binding. This constraint, which we and others have described for oligonucleotide loading into purified exosomes (Biscans *et al*., manuscript in press)^35–37^, provides a rationale for why hydrophobicity drives hsiRNA partitioning into distinct lipid transport pathways. VLDL and LDL carry highly-nonpolar saturated triglycerides and unsaturated cholesteryl esters, respectively, while HDL carries cholesterol and unsaturated phospholipids (relatively more polar). Our findings suggest that hsiRNAs equilibrate with circulating lipoproteins based on the solubility of the lipid conjugate in the lipoprotein core. Our results are corroborated by the observation that even when highly-lipophilic Chol-siRNA conjugates are exogenously loaded into HDL, they rapidly redistribute into LDL in serum^6^. We were surprised that none of our lipid-conjugated hsiRNAs showed significant affinity for albumin *in vivo*, despite our demonstration of robust albumin binding *in vitro*. Several recent studies have postulated that lipid-conjugated oligonucleotides leverage plasma albumin as an endogenous carrier^7,38^.

However, these studies used multivalent lipids, which may be better suited to access the multiple buried fatty acid binding sites in albumin.

Prior studies have established that cholesterol-conjugated siRNAs have 8-to 15-fold higher activity when pre-complexed into purified HDL than when injected alone, leading to a surge in the exploration of biomimetic lipoprotein nanoparticles as siRNA delivery vehicles^39–44^. One caveat to those reports is the use of unmodified or partially 2’-substituted siRNA scaffolds, which are inherently unstable in serum^45,46^. Our study demonstrates that using chemically-stabilized hsiRNAs removes the necessity for exogenous formulation to achieve potent mRNA silencing. A major advantage to using an siRNA delivery program that harnesses endogenous lipoproteins is the ease of oligonucleotide bioconjugation and biocompatibility. It also circumvents the typical challenges associated with using synthetic nanoparticles, including low loading efficiency, complex surface-modification processes, and systemic toxicity^47^. Although isolated and purified exosomes and HDL hold promise as non-immunogenic oligonucleotide carriers, they also face the dual challenges of low isolation and loading yields^48^. Finally, an important advantage of lipid-conjugation is that its impact on biodistribution is sequence-independent. Thus, this strategy should behave analogously for the delivery of other small RNA cargo, including miRNA replacements, antagomirs, or CRISPR components. Although lipid conjugation as a delivery platform has many strengths, it also has limitations, chief among them being difficulty in achieving cell type-specific delivery. This concern is being addressed in various forums, including the redirection of endogenous HDL and its therapeutic cargo via modification of surface apolipoprotein residues^49^.

Our findings suggest a revision of canonical design parameters for conjugated siRNAs. First, and of key importance, is an siRNA scaffold that contains substantial 2’-ribose and backbone modifications to improve nuclease stability^8^. Second, lipoprotein association will likely govern the systemic distribution of oligonucleotide conjugates (e.g. retinol or α-tocopherol) that are otherwise intended to be cell- or tissue-targeting^20^.

## Methods

### Oligonucleotide synthesis

Oligonucleotides were synthesized using standard and modified (2′-fluoro, 2′-*O*-methyl) phosphoramidite, solid-phase synthesis conditions using a MerMade 12 (BioAutomation) and Expedite ABI DNA/RNA synthesizer (ABI 8909). Oligonucleotides were removed from controlled pore glass (CPG), deprotected, and HPLC purified as described previously^8–10^. Ion exchange was performed on purified oligonucleotides using a Hi-Trap cation exchange column. The identity of oligonucleotides was established by LC-MS analysis (Waters Q-TOF premier). Relative degree of hydrophobicity of sense strands was assayed by reverse-phase HPLC (Waters Symmetric 3.5 µm, 4.6 x 75 mm column) using a 0-100% gradient over 15 minutes at 60°C with 0.1% TEAA in water (eluent A) and 100% acetonitrile (eluent B). Peaks were monitored at 260 nm.

### Oligonucleotide delivery

HeLa cells were plated in DMEM containing 6% FBS at 10,000 cells per well in 96-well tissue culture plates. hsiRNA was diluted to twice the final concentration in OptiMEM (Gibco), and mixed 1:1 with Lipofectamine RNAiMAX Transfection Reagent (Invitrogen) (final transfection reagent concentration = 0.3 μL/25 μL/well). 50 μL-diluted hsiRNA was added to 50 μL of cells, resulting in a final concentration of 3% FBS. Cells were incubated for 72 hours at 37°C and 5% CO2. For all *in vivo* studies, lipid-hsiRNAs were delivery without a transfection reagent.

### mRNA quantification

mRNA was quantified using the QuantiGene 2.0 Assay (Affymetrix) as described previously^8^. Briefly, cells were lysed in 250 μL-diluted lysis mixture with Proteinase K (Affymetrix) for 30 min at 55°C prior to mRNA quantification. Tissue punches (~5 mg) were homogenized in 300 μL of Homogenizing Buffer (Affymetrix) with proteinase K in 96-well plate format using a TissueLyser II (Qiagen). This method is described in detail in Coles *et al*^50^.

### Mouse studies

All animal procedures were approved by the University of Massachusetts Medical School Institutional Animal Care and Use Committee (IACUC, protocol number A-2411). Mice (FVB/NJ) were 6–10 weeks of age at the time of experiments. All animals were kept on a 12-h light/dark cycle in a pathogen-free facility with food and water *ad libitum*. For imaging and mRNA-silencing studies, mice were injected subcutaneously (SC, interscapular, between shoulders) or intravenously (IV, via tail vein) with 10-20 mg kg^−1^ of Cy3-labeled oligonucleotides (see figure captions for details). After 4 h-1 week, mice were deeply anesthetized with 0.1% Avertin and transcardially perfused with a 4% paraformaldehyde solution in phosphate-buffered saline, pH 7.2. Tissues were collected and incubated overnight at 4°C in either 10% formalin for subsequent imaging or RNALater (Ambion) for subsequent mRNA quantification. Tissues were processed as described previously^8^.

### Imaging and image quantification

All fluorescence images were acquired with a Leica DM5500 microscope fitted with a DFC365 FX fluorescence camera. After image acquisition, images were exported in 16-bit TIFF format from the Leica LAS X software and processed in Fiji v1.51n^51^ to quantify the kinetics of oligo uptake into cells. Briefly, the freehand tool was used to manually select individual cells and the mean fluorescence intensity was obtained by using the “Measure” function. Background fluorescence was corrected for by measuring the mean fluorescence of an area with no cells and subtracting this value from the cellular fluorescence intensity.

### Peptide nucleic acid (PNA) hybridization assay

Tissue accumulation of lipid-conjugated hsiRNAs was quantified as described previously^8,25^. Briefly, tissues were lysed in MasterPure tissue lysis solution (EpiCentre) in the presence of proteinase K (2 mg/ml) (Invitrogen) using a TissueLyser II (Qiagen) (~10 mg tissue per 100 μL lysis solution). Sodium dodecyl sulfate (SDS) was precipitated with KCl (3 M) and pelleted at 5000 x g. hsiRNAs presented in the cleared supernatant was hybridized to a Cy3-labeled PNA that was fully complementary to guide strand (PNABio) and injected on a DNAPac PA100 anion exchange column (Thermo Fisher). Cy3 fluorescence was monitored at 570 nm.

### Lipoprotein size exclusion chromatography

For lipoprotein profiling, mice were injected intravenously with 20 mg kg^−1^ of Cy3-labeled oligonucleotides. After 15 minutes, whole mouse blood (~500 µL) was collected in a sterile EDTA-coated tube following cheek incision with a lancet. Samples were spun at 10,000 RPM for 10 minutes at 4°C. 50 µL of serum was directly injected on Superose 6 Increase 10/300 size exclusion column (GE Healthcare). Oligonucleotide migration was monitored by Cy3 fluorescence at 570 nm, and lipoprotein protein content was monitored by absorbance at 280 nm.

### Statistical analysis

Data were analyzed using GraphPad Prism 6 software. IC_50_ curves were fitted using log(inhibitor) versus response––variable slope (four parameters). For Figure 4, statistics were calculated using one-way ANOVA with Tukey’s test for multiple comparisons, with significance calculated relative to saline control-injected animals. For Figure 5, statistics were calculated using an unpaired, two-tailed t-test.

### Data availability

The authors declare that all data supporting the findings of this study are available within the manuscript and supplementary information.

#### Acknowledgements

We would like to thank Emily Mohn and Darryl Conte for assistance with manuscript editing and all members of the Khvorova lab for their guidance. We thank Batuhan Yenilmez for help with preparing animal tissue for FACS analysis. This research was supported by 5F32NS095508-03 (to M.F.O.). Compound synthesis was supported by S10 OD 020012-01.

## Author Contributions

M.R.H., D.E., M.N., L.R., and A.B. synthesized all compounds. M.F.O., A.H.C., and B.M.D.C.G. performed all *in vivo* experiments and imaging. S.D. conducted the *in vitro* experiments. S.L. performed image quantification and statistical analysis. R.A.H. developed and conducted the PNA analysis. M.F.O., A.H.C., and A.K. came up with the concept. M.F.O. wrote the manuscript. A.K. edited the manuscript.

## References

1 Gao, S. et al. The effect of chemical modification and nanoparticle formulation on stability and biodistribution of siRNA in mice. Mol Ther 17, 1225–1233, doi:10.1038/mt.2009.91 (2009).

2 Juliano, R. L. The delivery of therapeutic oligonucleotides. Nucleic Acids Res 44, 6518–6548, doi:10.1093/nar/gkw236 (2016).

3 Fitzgerald, K. et al. A Highly Durable RNAi Therapeutic Inhibitor of PCSK9. N Engl J Med 376, 41–51, doi:10.1056/NEJMoa1609243 (2017).

4 Huang, Y. Preclinical and Clinical Advances of GalNAc-Decorated Nucleic Acid Therapeutics. Mol Ther Nucleic Acids 6, 116–132, doi:10.1016/j.omtn.2016.12.003 (2017).

5 Nishina, K. et al. Efficient In Vivo Delivery of siRNA to the Liver by Conjugation of alpha-Tocopherol. Mol Ther 16, 734–740, doi:10.1038/mt.2008.14 (2008).

6 Wolfrum, C. et al. Mechanisms and optimization of in vivo delivery of lipophilic siRNAs. Nat Biotechnol 25, 1149–1157, doi:10.1038/nbt1339 (2007).

7 Sarett, S. M. et al. Lipophilic siRNA targets albumin in situ and promotes bioavailability, tumor penetration, and carrier-free gene silencing. Proc Natl Acad Sci U S A 114, E6490–E6497, doi:10.1073/pnas.1621240114 (2017).

8 Hassler, M. R. et al. Comparison of partially and fully chemically-modified siRNA in conjugate-mediated delivery in vivo. Nucleic Acids Res, doi:10.1093/nar/gky037 (2018).

9 Alterman, J. F. et al. Hydrophobically Modified siRNAs Silence Huntingtin mRNA in Primary Neurons and Mouse Brain. Mol Ther Nucleic Acids 4, e266, doi:10.1038/mtna.2015.38 (2015).

10 Nikan, M. et al. Docosahexaenoic Acid Conjugation Enhances Distribution and Safety of siRNA upon Local Administration in Mouse Brain. Mol Ther Nucleic Acids 5, e344, doi:10.1038/mtna.2016.50 (2016).

11 Nikan, M. et al. Synthesis and Evaluation of Parenchymal Retention and Efficacy of a Metabolically Stable O-Phosphocholine-N-docosahexaenoyl-l-serine siRNA Conjugate in Mouse Brain. Bioconjug Chem 28, 1758–1766, doi:10.1021/acs.bioconjchem.7b00226 (2017).

12 Allerson, C. R. et al. Fully 2’-modified oligonucleotide duplexes with improved in vitro potency and stability compared to unmodified small interfering RNA. J Med Chem 48, 901–904, doi:10.1021/jm049167j (2005).

13 Haraszti, R. A. et al. 5-Vinylphosphonate improves tissue accumulation and efficacy of conjugated siRNAs in vivo. Nucleic Acids Res, doi:10.1093/nar/gkx507 (2017).

14 Geary, R. S., Norris, D., Yu, R. & Bennett C. F. Pharmacokinetics, biodistribution and cell uptake of antisense oligonucleotides. Adv Drug Deliv Rev 87, 46–51, doi:10.1016/j.addr.2015.01.008 (2015).

15 Byrne, M. et al. Novel hydrophobically modified asymmetric RNAi compounds (sd-rxRNA) demonstrate robust efficacy in the eye. J Ocul Pharmacol Ther 29, 855–864, doi:10.1089/jop.2013.0148 (2013).

16 Godinho, B. et al. Pharmacokinetic Profiling of Conjugated Therapeutic Oligonucleotides: A High-Throughput Method Based Upon Serial Blood Microsampling Coupled to Peptide Nucleic Acid Hybridization Assay. Nucleic Acid Ther, doi:10.1089/nat.2017.0690 (2017).

17 Ghosh, N., Mondal, R. & Mukherjee S. Hydrophobicity is the governing factor in the interaction of human serum albumin with bile salts. Langmuir 31, 1095–1104, doi:10.1021/la504270a (2015).

18 van der Sluis, R. J., Van Eck, M. & Hoekstra, M. Adrenocortical LDL receptor function negatively influences glucocorticoid output. J Endocrinol 226, 145–154, doi:10.1530/JOE-15-0023 (2015).

19 Donner, A. J. et al. Co-Administration of an Excipient Oligonucleotide Helps Delineate Pathways of Productive and Nonproductive Uptake of Phosphorothioate Antisense Oligonucleotides in the Liver. Nucleic Acid Ther 27, 209–220, doi:10.1089/nat.2017.0662 (2017).

20 Murakami, M. et al. Enteral siRNA delivery technique for therapeutic gene silencing in the liver via the lymphatic route. Sci Rep 5, 17035, doi:10.1038/srep17035 (2015).

21 Schmidt, K. et al. Characterizing the effect of GalNAc and phosphorothioate backbone on binding of antisense oligonucleotides to the asialoglycoprotein receptor. Nucleic Acids Res 45, 2294–2306, doi:10.1093/nar/gkx060 (2017).

22 Tanowitz, M. et al. Asialoglycoprotein receptor 1 mediates productive uptake of N-acetylgalactosamine-conjugated and unconjugated phosphorothioate antisense oligonucleotides into liver hepatocytes. Nucleic Acids Res 45, 12388–12400, doi:10.1093/nar/gkx960 (2017).

23 Molitoris, B. A. et al. siRNA targeted to p53 attenuates ischemic and cisplatin-induced acute kidney injury. J Am Soc Nephrol 20, 1754–1764, doi:10.1681/ASN.2008111204 (2009).

24 Thompson, J. D. et al. Toxicological and pharmacokinetic properties of chemically modified siRNAs targeting p53 RNA following intravenous administration. Nucleic Acid Ther 22, 255–264, doi:10.1089/nat.2012.0371 (2012).

25 Haraszti, R. A. et al. 5-Vinylphosphonate improves tissue accumulation and efficacy of conjugated siRNAs in vivo. Nucleic Acids Res 45, 7581–7592, doi:10.1093/nar/gkx507 (2017).

26 Wagner, J. et al. Characterization of levels and cellular transfer of circulating lipoprotein-bound microRNAs. Arterioscler Thromb Vasc Biol 33, 1392–1400, doi:10.1161/ATVBAHA.112.300741 (2013).

27 Vickers, K. C., Palmisano, B. T., Shoucri, B. M., Shamburek, R. D. & Remaley, A. T. MicroRNAs are transported in plasma and delivered to recipient cells by high-density lipoproteins. Nat Cell Biol 13, 423–433, doi:10.1038/ncb2210 (2011).

28 Tabet, F. et al. HDL-transferred microRNA-223 regulates ICAM-1 expression in endothelial cells. Nat Commun 5, 3292, doi:10.1038/ncomms4292 (2014).

29 Ono, K. et al. MicroRNAs and High-Density Lipoprotein Cholesterol Metabolism. Int Heart J 56, 365–371, doi:10.1536/ihj.15-019 (2015).

30 Mengistu, D. H., Bohinc, K. & May, S. Binding of DNA to zwitterionic lipid layers mediated by divalent cations. J Phys Chem B 113, 12277–12282, doi:10.1021/jp904986j (2009).

31 Janas, T., Janas, T. & Yarus, M. Specific RNA binding to ordered phospholipid bilayers. Nucleic Acids Res 34, 2128–2136, doi:10.1093/nar/gkl220 (2006).

32 Gromelski, S. & Brezesinski, G. DNA condensation and interaction with zwitterionic phospholipids mediated by divalent cations. Langmuir 22, 6293–6301, doi:10.1021/la0531796 (2006).

33 Lu, D. & Rhodes, D. G. Binding of phosphorothioate oligonucleotides to zwitterionic liposomes. Biochim Biophys Acta 1563, 45–52 (2002).

34 Khvorova, A., Kwak, Y. G., Tamkun, M., Majerfeld, I. & Yarus, M. RNAs that bind and change the permeability of phospholipid membranes. Proc Natl Acad Sci U S A 96, 10649–10654 (1999).

35 Didiot, M. C. et al. Exosome-mediated Delivery of Hydrophobically Modified siRNA for Huntingtin mRNA Silencing. Mol Ther 24, 1836–1847, doi:10.1038/mt.2016.126 (2016).

36 Haraszti, R. A., Coles, A., Aronin, N., Khvorova, A. & Didiot, M. C. Loading of Extracellular Vesicles with Chemically Stabilized Hydrophobic siRNAs for the Treatment of Disease in the Central Nervous System. Bio Protoc 7, doi:10.21769/BioProtoc.2338 (2017).

37 O’Loughlin, A. J. et al. Functional Delivery of Lipid-Conjugated siRNA by Extracellular Vesicles. Mol Ther 25, 1580–1587, doi:10.1016/j.ymthe.2017.03.021 (2017).

38 Lacroix, A., Edwardson, T. G. W., Hancock, M. A., Dore, M. D. & Sleiman, H. F. Development of DNA Nanostructures for High-Affinity Binding to Human Serum Albumin. J Am Chem Soc 139, 7355–7362, doi:10.1021/jacs.7b02917 (2017).

39 Yang, M. et al. Efficient cytosolic delivery of siRNA using HDL-mimicking nanoparticles. Small 7, 568–573, doi:10.1002/smll.201001589 (2011).

40 Zhang, Z. et al. HDL-mimicking peptide-lipid nanoparticles with improved tumor targeting. Small 6, 430–437, doi:10.1002/smll.200901515 (2010).

41 Lin, Q. et al. Efficient systemic delivery of siRNA by using high-density lipoprotein-mimicking peptide lipid nanoparticles. Nanomedicine (Lond) 7, 1813–1825, doi:10.2217/nnm.12.73 (2012).

42 Nakayama, T. et al. Harnessing a physiologic mechanism for siRNA delivery with mimetic lipoprotein particles. Mol Ther 20, 1582–1589, doi:10.1038/mt.2012.33 (2012).

43 Kuwahara, H. et al. Efficient in vivo delivery of siRNA into brain capillary endothelial cells along with endogenous lipoprotein. Mol Ther 19, 2213–2221, doi:10.1038/mt.2011.186 (2011).

44 Huang, J. L. et al. Lipoprotein-biomimetic nanostructure enables efficient targeting delivery of siRNA to Ras-activated glioblastoma cells via macropinocytosis. Nat Commun 8, 15144, doi:10.1038/ncomms15144 (2017).

45 Ding, Y. et al. A biomimetic nanovector-mediated targeted cholesterol-conjugated siRNA delivery for tumor gene therapy. Biomaterials 33, 8893–8905, doi:10.1016/j.biomaterials.2012.08.057 (2012).

46 Kennedy, S., Wang, D. & Ruvkun, G. A conserved siRNA-degrading RNase negatively regulates RNA interference in C. elegans. Nature 427, 645–649, doi:10.1038/nature02302 (2004).

47 Miele, E. et al. Nanoparticle-based delivery of small interfering RNA: challenges for cancer therapy. Int J Nanomedicine 7, 3637–3657, doi:10.2147/IJN.S23696 (2012).

48 Kuai, R., Li, D., Chen, Y. E., Moon, J. J. & Schwendeman, A. High-Density Lipoproteins: Nature’s Multifunctional Nanoparticles. ACS Nano 10, 3015–3041, doi:10.1021/acsnano.5b07522 (2016).

49 Ding, Y. et al. Rerouting Native HDL to Predetermined Receptors for Improved Tumor-Targeted Gene Silencing Therapy. ACS Appl Mater Interfaces 9, 30488–30501, doi:10.1021/acsami.7b10047 (2017).

50 Coles, A. H. et al. A High-Throughput Method for Direct Detection of Therapeutic Oligonucleotide-Induced Gene Silencing In Vivo. Nucleic Acid Ther 26, 86–92, doi:10.1089/nat.2015.0578 (2016).

51 Schindelin, J. et al. Fiji: an open-source platform for biological-image analysis. Nat Methods 9, 676–682, doi:10.1038/nmeth.2019 (2012).

