## Supplementary Materials for "Hydrophobicity drives the systemic distribution of lipid-conjugated siRNAs via lipid transport pathways"

[*]These authors contributed equally to this work
[†]Present address: Ionis Pharmaceuticals, Carlsbad, CA
[‡]Co-corresponding authors

*Table of Contents*

| | |
|---|---|
| Supplementary Materials and Methods | S2 |
| **Table S1.** Oligonucleotide sequences and chemical modification patterns | S3 |
| **Table S2.** Predicted lipid partition coefficients | S3 |
| **Figure S1.** Lipid-hsiRNAs interact with albumin | S4 |
| **Figure S2.** Lipoprotein chromatography and cholesterol content in male and female plasma | S5 |
| **Table S3.** Summary of peak integrations from lipoprotein binding profiles | S6 |
| **Figure S3.** FACS profiling of lipid-hsiRNA accumulation in liver | S7 |
| **Figure S4.** mRNA silencing following administration of non-targeting controls | S8 |
| **Figure S5.** Lipid-hsiRNA treatment does not induce any changes in blood chemistry | S8 |

### Supplementary Materials and Methods

#### Gel shift assay

Lipid-hsiRNAs (5 μM) were incubated with increasing bovine serum albumin (0-200 μM or 0, 2, 5, 8, and 10 g/L) in 20 mM Tris pH 7.4. Samples was incubated for 17 h on an orbital shaker (100 rpm) at room temperature and subsequently analyzed by 6% native TBE EMSA. Bands were quantified using ImageJ software (ImageJ 1.50i) and affinity constants were calculated for DCA-hsiRNA and Chol-hsiRNA by determining the fraction of lipid-hsiRNA bound in each lane.

#### Fluorescence associated cell sorting (FACS)

Animals were injected subcutaneously (SC, interscapular, between shoulders) with 10 mg $kg^{-1}$ of Cy3-labeled oligonucleotides (n = 3). After 1 week, mice were deeply anesthetized with 0.1% Avertin and livers were dissected and prepared for downstream analysis exactly as described previously[1]. hsiRNA, DCA-hsiRNA, and DHA-hsiRNA intracellular accumulation was measured in CD31+ endothelial cells and F4/80+ Kupffer cells by flow cytometry using an LSR II cell analyzer (Becton-Dickenson). Fluorescence-activated cell sorter analyses were prepared by FlowJo software (Tree Star Inc., Ashland, OR).

#### Toxicity

At the termination of the experiment described in Figure 4 (1 week after administration of test compounds), mouse blood (~500 μL) was collected in a sterile lithium heparin-coated tube following cheek incision with a lancet (n = 6). Samples were spun at 10,000 RPM for 10 minutes at 4°C to isolate serum. The UMMS National Mouse Metabolic Phenotyping Core analyzed the samples for the following parameters: blood urea nitrogen (BUN); alanine aminotransferase (ALT); alkaline phosphatase (ALP); albumin (ALB); amylase (AMY); calcium ($Ca^{2+}$); creatinine (CRE); globulin (GLOB); glucose (GLU); phosphorus (PHOS); potassium (K+); sodium (Na+); total bilirubin (TBIL), using an Abaxis comprehensive rotor.

#### Serum cholesterol measurement

For serum cholesterol quantification, whole mouse blood (~500 μL) was collected in a sterile EDTA-coated tube following cheek incision with a lancet from both male and female mice (n = 2). Samples were spun at 10,000 RPM for 10 minutes at 4°C. 50 μL of serum was directly injected on Superose 6 Increase 10/300 size exclusion column (GE Healthcare). Oligonucleotide migration was monitored by Cy3 fluorescence at 570 nm, and lipoprotein protein content was monitored by absorbance at 280 nm. Fractions were collected and analyzed for cholesterol content using the HDL and LDL Cholesterol Assay Kit (Abcam) according to the manufacturer's instructions.

| siRNA ID | Gene | Accession Number | Targeting Position | Strand | Chemical Modification Pattern | Conjugate |
|---|---|---|---|---|---|---|
| hsiRNA $^{Ppib}$ | PPIB | NM_009693.2 | 437 | S | fC#mA#fA.mA.fU.mU.fC.mC.fA.mU.fC. mG.fU#mG#fA | |
| Chol-hsiRNA $^{Ppib}$ | PPIB | NM_009693.2 | 437 | S | fC#mA#fA.mA.fU.mU.fC.mC.fA.mU.fC. mG.fU#mG#fA | 3'-Cholesterol |
| LCA-hsiRNA $^{Ppib}$ | PPIB | NM_009693.2 | 437 | S | fC#mA#fA.mA.fU.mU.fC.mC.fA.mU.fC. mG.fU#mG#fA | 3'-Lithocholic acid |
| DHA-hsiRNA $^{Ppib}$ | PPIB | NM_009693.2 | 437 | S | fC#mA#fA.mA.fU.mU.fC.mC.fA.mU.fC. mG.fU#mG#fA | 3'-Docosahexaenoic acid |
| DCA-hsiRNA $^{Ppib}$ | PPIB | NM_009693.2 | 437 | S | fC#mA#fA.mA.fU.mU.fC.mC.fA.mU.fC. mG.fU#mG#fA | 3'-Docosanoic acid |
| hsiRNA $^{Ppib}$ | PPIB | NM_009693.2 | 437 | AS | VPmU#fU#mA.fA.mU.fC.mU.fC.mU.fU. mU.fA.mC#fU#mG#fA#mU#fA#mU#fA | |
| Cy3-hsiRNA $^{Ppib}$ | PPIB | NM_009693.2 | 437 | S | fC#mA#fA.mA.fU.mU.fC.mC.fA.mU.fC. mG.fU#mG#fA | |
| Cy3-Chol-hsiRNA $^{Ppib}$ | PPIB | NM_009693.2 | 437 | S | fC#mA#fA.mA.fU.mU.fC.mC.fA.mU.fC. mG.fU#mG#fA | 3'-Cholesterol; 5'-Cy3 |
| Cy3-LCA-hsiRNA $^{Ppib}$ | PPIB | NM_009693.2 | 437 | S | fC#mA#fA.mA.fU.mU.fC.mC.fA.mU.fC. mG.fU#mG#fA | 3'-Lithocholic acid; 5'-Cy3 |
| Cy3-DHA-hsiRNA $^{Ppib}$ | PPIB | NM_009693.2 | 437 | S | fC#mA#fA.mA.fU.mU.fC.mC.fA.mU.fC. mG.fU#mG#fA | 3'-Docosahexaenoic acid; 5'-Cy3 |
| Cy3-DCA-hsiRNA $^{Ppib}$ | PPIB | NM_009693.2 | 437 | S | fC#mA#fA.mA.fU.mU.fC.mC.fA.mU.fC. mG.fU#mG#fA | 3'-Docosanoic acid; 5'-Cy3 |
| hsiRNA $^{NTC}$ | NTC | -- | -- | S | fU#mG#fA.mC.fA.mA.fA.mU.fA.mC.fG. mA.fU#mU#fA | |
| Chol-hsiRNA $^{NTC}$ | NTC | -- | -- | S | fU#mG#fA.mC.fA.mA.fA.mU.fA.mC.fG. mA.fU#mU#fA | 3'-Cholesterol |
| LCA-hsiRNA $^{NTC}$ | NTC | -- | -- | S | fU#mG#fA.mC.fA.mA.fA.mU.fA.mC.fG. mA.fU#mU#fA | 3'-Lithocholic acid |
| DHA-hsiRNA $^{NTC}$ | NTC | -- | -- | S | fU#mG#fA.mC.fA.mA.fA.mU.fA.mC.fG. mA.fU#mU#fA | 3'-Docosahexaenoic acid |
| DCA-hsiRNA $^{NTC}$ | NTC | -- | -- | S | fU#mG#fA.mC.fA.mA.fA.mU.fA.mC.fG. mA.fU#mU#fA | 3'-Docosanoic acid |
| hsiRNA $^{NTC}$ | NTC | -- | -- | AS | VPmU#fA#mA.fU.mC.fG.mU.fA.mU.fU. mU.fG.mU#fC#mA#fA#mU#fC#mA#fU | |
| Blunt Cy3-DHA-hsiRNA $^{Ppib}$ | PPIB | NM_009693.2 | 437 | S | mA#fA#mC.fA.mG.fC.mA.fA.mA.fU.mU .fC.mC.fA.mU.fC.mG.fU#mG#fA | 3'-Docosahexaenoic acid; 5'-Cy3 |
| Blunt Cy3-DCA-hsiRNA $^{Ppib}$ | PPIB | NM_009693.2 | 437 | S | mA#fA#mC.fA.mG.fC.mA.fA.mA.fU.mU .fC.mC.fA.mU.fC.mG.fU#mG#fA | 3'-Docosanoic acid; 5'-Cy3 |
| Blunt Cy3-GalNAc-hsiRNA $^{Pp}$ | PPIB | NM_009693.2 | 437 | S | mA#fA#mC.fA.mG.fC.mA.fA.mA.fU.mU .fC.mC.fA.mU.fC.mG.fU#mG#fA | 3'-N-acetylgalactosamine; 5'-Cy3 |
| GalNAc-hsiRNA $^{Ppib}$ | PPIB | NM_009693.2 | 437 | S | fC#mA#fA.mA.fU.mU.fC.mC.fA.mU.fC. mG.fU#mG#fA | 3'-N-acetylgalactosamine; 5'-Cy3 |
| Blunt hsiRNA $^{Ppib}$ | PPIB | NM_009693.2 | 437 | AS | VPmU#fU#mA.fA.mU.fC.mU.fC.mU.fU. mU.fA.mC.fU.mG.fA.mU.fA#mU#fA | |

**Table S1. Modified oligonucleotide sequences.** Chemical modifications are abbreviated as follows, where "X" corresponds to A, U, G, or C: fX (2'-fluoro), mX (2'-O-methyl), VP (5'-vinylphosphonate), '#' (phosphorothioate backbone modification), '.' (phosphodiester backbone). Oligonucleotide synthesis procedures are described in the Materials and Methods.

| Molecule | Predicted LogP |
|---|---|
| Cholesterol | 7.62 |
| Docosanoic acid | 9.13 |
| Docosahexaenoic acid | 5.68 |
| Lithocholic acid | 5.16 |

**Table S2. Predicted partition coefficients for each lipid.** logP values determined using the Molinspiration Interactive logP calculator (Molinspiration Property Calculation Service, www.molinspiration.com).

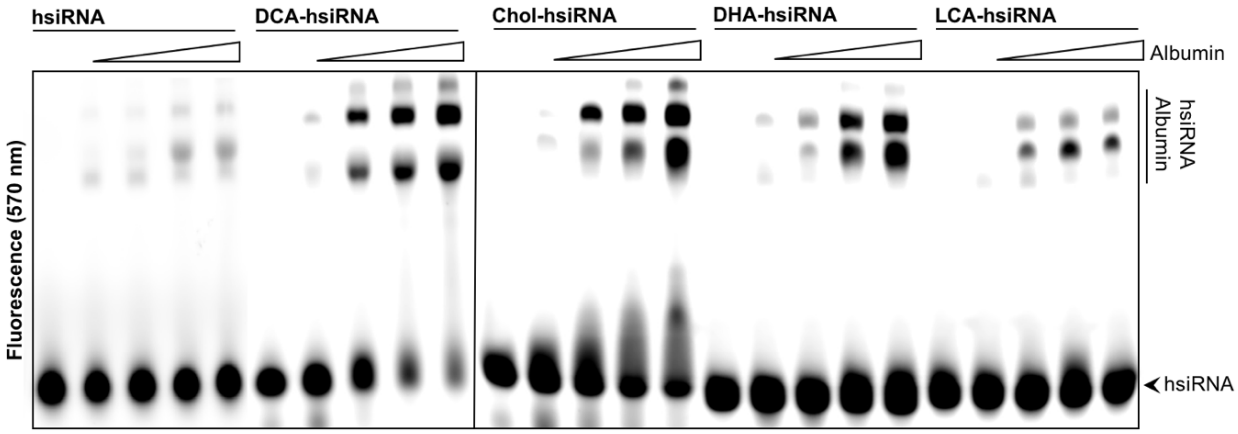


**Figure S1. Lipid-hsiRNAs interact with albumin.** Gel shift assay between lipid-hsiRNAs (5 $\mu$M) in the presence of increasing bovine serum albumin (0-200 $\mu$M, 0, 2, 5, 8, and 10 g/L).

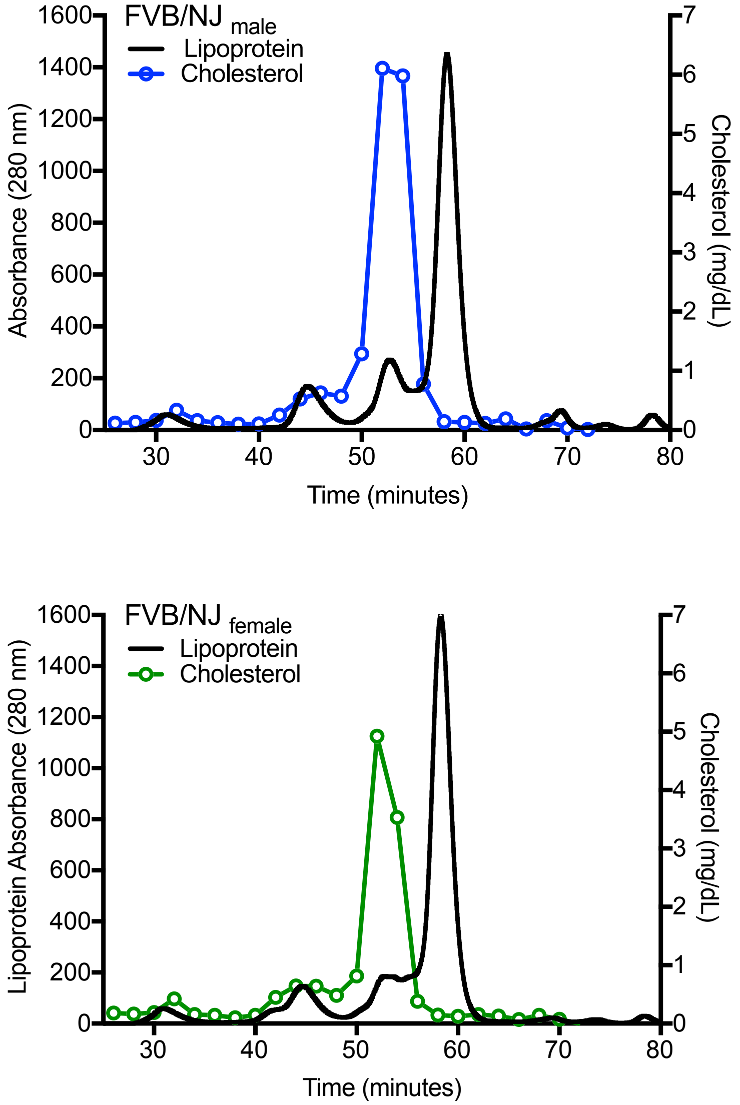


**Figure S2. Lipoprotein and cholesterol profiling in male and female mouse plasma.** Mouse serum lipoprotein and cholesterol distribution following size exclusion chromatography (SEC) from male (upper panel) or female (lower panel) wild-type animals. Black lines: serum proteins; blue/green lines: cholesterol.

| | VLDL | LDL | HDL | Albumin | Unbound | Conjugate |
|---|---|---|---|---|---|---|
| hsiRNA | 0.0 | 0.0 | 0.0 | 0.0 | 99.5 | |
| LCA-hsiRNA | 12.2 | 5.3 | 50.4 | 32.2 | 0.0 | Lithocholic acid |
| PC-DHA-hsiRNA | 2.9 | 5.0 | 60.5 | 31.7 | 0.0 | Phosphocholine-docosahexaenoic acid |
| EPA-hsiRNA | 9.3 | 6.1 | 61.6 | 23.0 | 0.0 | Eicosapentanoic acid |
| DHA-hsiRNA | 1.0 | 3.7 | 80.5 | 14.9 | 0.0 | Docosahexaenoic acid |
| RA-hsiRNA | 1.8 | 11.7 | 86.4 | 0.0 | 0.0 | Retionic acid |
| PC-DCA-hsiRNA | 2.7 | 92.2 | 4.9 | 0.0 | 0.0 | Phosphocholine-docosanoic acid |
| DCA-hsiRNA | 19.1 | 65.0 | 11.1 | 0.0 | 0.0 | Docosanoic acid |
| Chol-hsiRNA | 7.3 | 82.2 | 10.5 | 0.0 | 0.0 | Cholesterol |
| GM1-hsiRNA | 0.0 | 93.8 | 6.2 | 0.0 | 0.0 | GM1 ganglioside |

**Table S3. Lipid-hsiRNAs partition into distinct LDL and HDL binding classes based on hydrophobicity.** Lipoprotein binding profiles were collected for a series of lipid-hsiRNAs (as described in Figure 3). Peaks were auto-selected and integrated using Agilent ChemStation software.

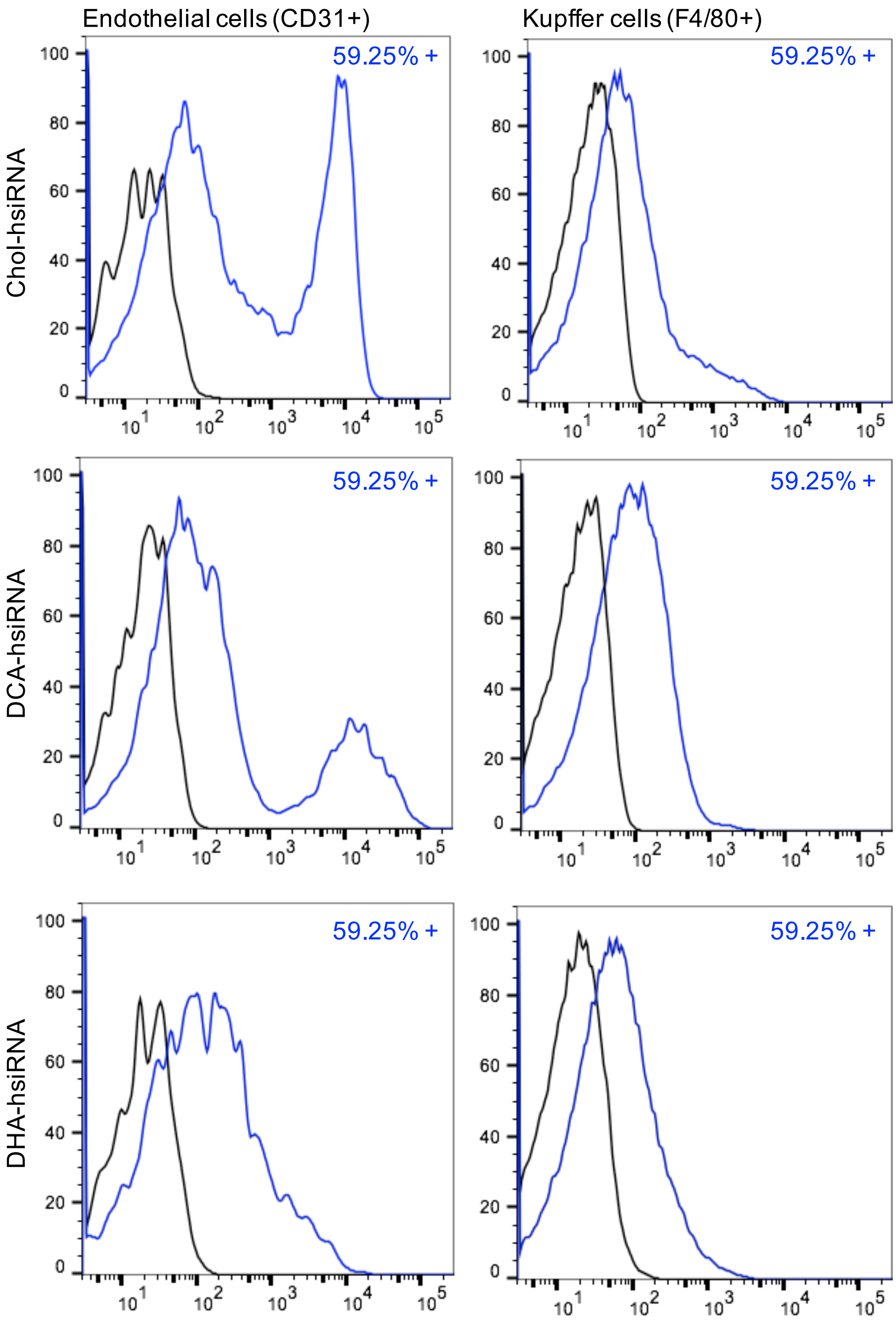


**Figure S3. FACS profiling of lipid-hsiRNA accumulation in liver.** hsiRNA, DCA-hsiRNA, and DHA-hsiRNA accumulation in CD31+ endothelial cells and F4/80+ Kupffer cells (stellate macrophages).

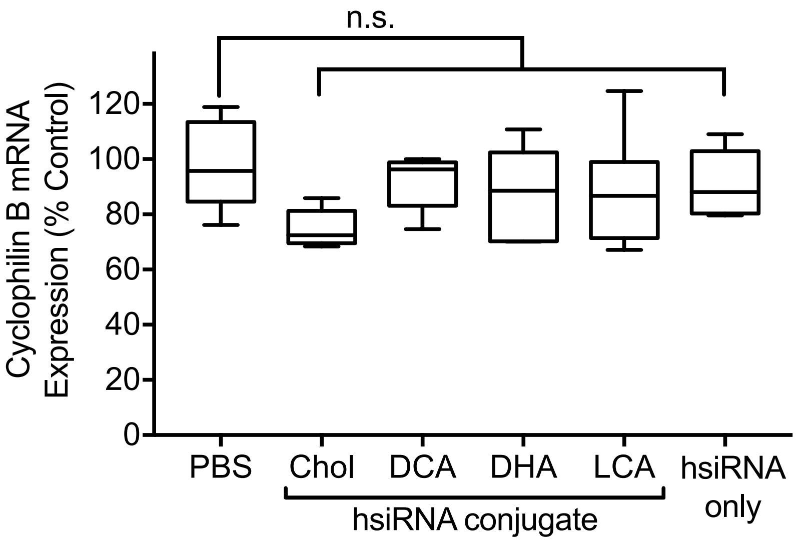


**Figure S4. mRNA silencing following administration of non-targeting controls.** Quantification of *Ppib* silencing by non-targeting control lipid-hsiRNAs in the liver. *Ppib* mRNA levels were measured with QuantiGene 2.0 (Affymetrix) assay and normalized to a housekeeping gene, *Hprt*. All data presented as percent of saline-treated control. All error bars represent mean ± SD. n.s. non-significant; *P < 0.05; **P < 0.01; ***P < 0.001; ****P < 0.0001 as calculated by one-way ANOVA with Tukey's test for multiple comparisons.

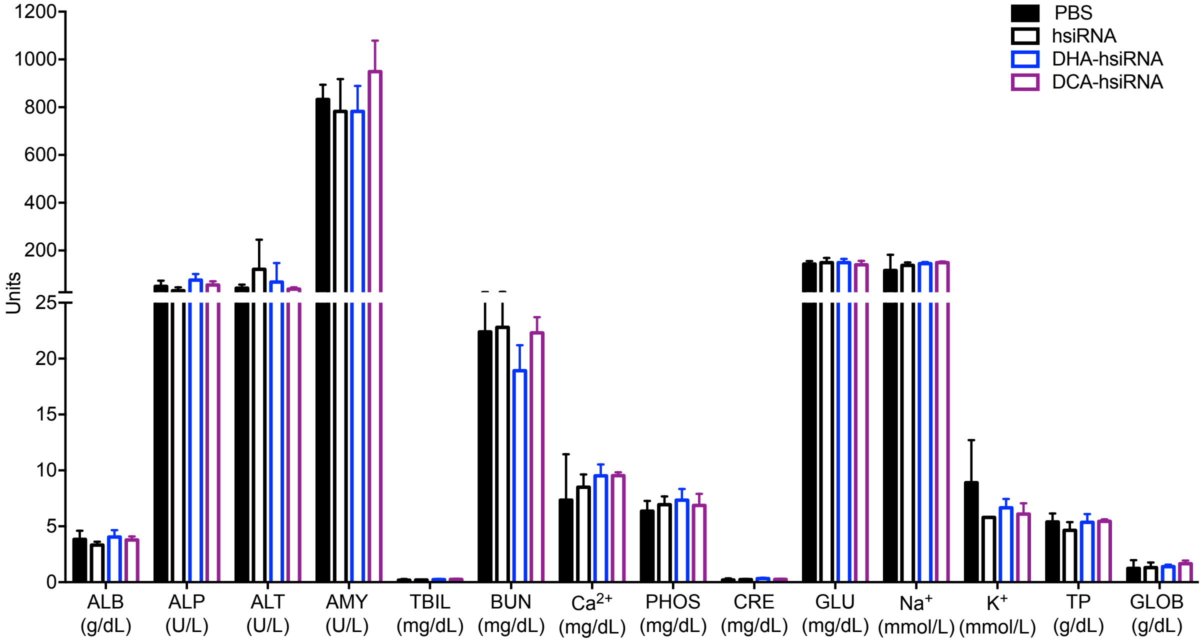


**Figure S5. Lipid-hsiRNA treatment does not induce any changes in blood chemistry.** A comprehensive panel of serum toxicological markers were measured one week after administration of test compounds (20 mg $kg^{-1}$, SC). No significant differences were observed between saline and hsiRNA-treated animals.

**References**

1 Severgnini, M. *et al*. A rapid two-step method for isolation of functional primary mouse hepatocytes: cell characterization and asialoglycoprotein receptor based assay development. *Cytotechnology* **64**, 187-195, doi:10.1007/s10616-011-9407-0 (2012).
